# Toxinome - The Bacterial Protein Toxin Database

**DOI:** 10.1101/2023.08.12.553073

**Authors:** Aleks Danov, Ofir Segev, Avi Bograd, Yedidya Ben Eliyahu, Noam Dotan, Tommy Kaplan, Asaf Levy

## Abstract

Protein toxins are key molecular weapons in biology that are used to attack neighboring cells. Bacteria use protein toxins to kill or inhibit growth of prokaryotic and eukaryotic cells using various modes of action that target essential cellular components. The toxins are responsible for shaping microbiomes in different habitats, for abortive phage infection, and for severe infectious diseases of animals and plants. Although several toxin databases have been developed, each one is devoted to a specific toxin family and they encompass a relatively small number of toxins. Antimicrobial toxins are often accompanied by antitoxins (or immunity proteins) that neutralize the cognate toxins. Here, we combined toxins and antitoxins from many resources and created Toxinome, a comprehensive and updated bacterial protein toxin database. Toxinome includes a total of 1,483,028 toxins and 491,345 antitoxins encoded in 59,475 bacterial genomes across the tree of life. We identified a depletion of toxin and antitoxin genes in bacteria that dwell in extreme temperatures. We defined 5,161 unique Toxin Islands within phylogenetically diverse bacterial genomes, which are loci dense in toxin and antitoxin genes. By focusing on the unannotated genes within these islands, we characterized a number of these genes as toxins or antitoxins. Finally, we developed an interactive Toxinome website (http://toxinome.pythonanywhere.com) that allows searching and downloading of our database. The Toxinome resource will be useful to the large research community interested in bacterial toxins and can guide toxin discovery and function elucidation, and infectious disease diagnosis and treatment.

**Importance:** Microbes use protein toxins as important tools to attack neighboring cells, microbial or eukaryotic, and for self-killing when attacked by viruses. These toxins work by different mechanisms to inhibit cell growth or kill cells. Microbes also use antitoxin proteins to neutralize the toxin activities. Here, we developed a comprehensive database called Toxinome of nearly two million toxin and antitoxins that are encoded in 59,475 bacterial genomes. We described the distribution of bacterial toxins and identified that they are depleted from bacteria that live in hot and cold temperatures. We find 5,161 cases in which toxins and antitoxins are densely clustered in bacterial genomes and termed these areas “Toxin Islands”. The Toxinome database is a useful resource for anyone interested in toxin biology and evolution, and it can guide discovery of new toxins.

## Introduction

Bacteria employ protein toxins to harm surrounding eukaryotic or prokaryotic cells. There is a large variety of bacterial protein toxins. These can be coarsely divided into several large classes: (a) effectors proteins that are translocated into target cells via different membrane-bound or extracellular bacterial secretion systems, such as the Type VI secretion system, contact-dependent inhibition systems, Tc toxins, or the extracellular contractile injection system (1–7), (b) toxins that are released from the attacking microbe and enter the target cell through specific receptors or through insertion into target cell membranes, e.g. bacteriocins, AB toxins or MARTX toxins (8–13), and (c) toxins that inhibit self growth of the producing cell in response to phage infection or antibiotic persistence, such as toxin-antitoxin systems (14–16). Toxin-antitoxin systems are currently divided into eight types and are implicated in impairment of DNA replication, translation, cell envelope, and cytoskeleton integrity, and they can induce metabolic stress (14, 17). Most of these toxins are studied in model bacterial genomes that represent only a minor portion of the over 600,000 bacterial genomes that have been sequenced in recent years (18). A more comprehensive view of all bacterial protein toxins will facilitate general understanding of toxin evolution, and inference of correlations between different toxin families and the genome organization of toxins. In addition, since in bacteria gene proximity often implies a functional relationship between neighboring genes (e.g. genes that are co-transcribed as part of an operon), an extensive toxin database can enable discovery of new toxins and toxin-associated genes based on their genomic neighborhood. Toxin-associated genes located next to toxins can be involved in toxin production, maturation or secretion, in immunity against the toxin or in lateral transfer of the toxin gene between bacteria.

There have been previous efforts to construct protein toxin databases. However, these databases often focus on one toxin class or they do not cover a large collection of bacterial protein toxins. The Toxin Exposome Database (T3DB) focuses mostly on drugs, industrial toxins, and pollutants but contains only few bacterial protein toxins (19). The Secret6 database is a comprehensive database of only microbial T6SS effector proteins, which are mostly serving as antimicrobial toxins (20). Similarly, TADB is an excellent resource for type 2 toxin-antitoxin systems, mostly used for self growth inhibition of bacteria to achieve plasmid addiction or dormancy (21). Bactibase and BAGEL4 focus on bacterial bacteriocins (22–24). DBETH includes exotoxins from human pathogens (22) and DBAASP is an extensive updated database of mostly synthetic antimicrobial peptides (25). A recent database, prokaryotic antimicrobial toxin database (PAT), is more inclusive than the aforementioned databases and contains 441 bacterial and archaeal toxins from seven classes, 64% of which are bacteriocins and T6SS effectors of proteobacteria (26). PAT also contains 6,064 predicted antimicrobial toxins in prokaryotic reference genomes. Notwithstanding, the vast majority of microbial protein toxins are currently not present in any toxin database except being part of general large protein databases, such as Uniprot. The microbiology research community therefore lacks an integrated updated database which focuses on microbial toxins from various classes and their mapping to tens of thousands microbial genomes. There are also recent reports that show that the same bacterial toxins can serve in one organism as part of self-inhibiting toxin-antitoxin modules, and in another organism they evolved into toxin effectors that are being injected into target cells (5, 27, 28), and hence there is little benefit in separating toxins into different databases based on their specific biological function. Moreover, numerous toxin proteins are likely encoded in the tens of thousands of bacterial genomes, many of which are poorly studied non-model or non-pathogenic bacteria. These toxin genes are currently not reported anywhere and they can aid in studying toxin function and evolution.

We developed “Toxinome”: The Bacterial Protein Toxin Database, which includes 1,483,028 toxins, and 491,345 immunity genes, from many toxin families that are encoded in 59,475 bacterial genomes. We developed an interactive interface which allows online querying and full downloading of the Toxinome database. Our analysis indicated that toxins are depleted from bacteria that dwell in extreme temperatures. Finally, using the Toxinome database we defined a large collection of 5,161 genomic regions that are highly rich in toxin and antitoxin genes which we define as genomic Toxin Islands. The Toxin Islands can be used to identify novel toxin and toxin-associated genes that are currently functionally unannotated. The resource we developed will be valuable to the large research community that studies different classes of bacterial toxins. It can be used for toxin and antitoxin annotations of genomes of interest, and it can be mined in an effort to discover new antimicrobial proteins, anticancer proteins, or biopesticides of clinical, biotechnological, and environmental importance.

## Materials and Methods

### Assembly of a toxin protein database

We initiated Toxinome by combining toxic proteins gathered from four databases: Secret6 (20), BactiBase (20, 23), TADB (21), Bagel (24), that were downloaded in October 2020. In addition, we collected a large number of toxin proteins from the UniProt database (29) using keyword searches (e.g., ‘toxic’, ‘toxin’). At this point we removed proteins with keywords such as ‘anti-toxin’ and ‘immunity’. These antitoxins were later added based on the Pfam database (see below). In total, 175,573 toxin protein sequences were included in the combined database. We clustered the 175,573 collected toxins using the CD-HIT 4.8.1 software (threshold = 0.7, alignment coverage = 0.92) (30). The proteins were clustered into 70,867 groups. We then mapped the toxin genes into microbial genomes downloaded from Integrated Microbial Genomic (IMG) database (31); the representative genes of all clusters were mapped to 1,078,446 genes from 59,475 genomes (Supplementary Table 1). The mapping was performed using DIAMOND 2.0.4.142 software (alignment coverage = 0.95, e-value = 0.05, threshold identity > 30%) (Figure 1) (32). To increase the number of toxins and immunity proteins into our database we used protein domain information. We added 219 and 94 toxin and antitoxin domains, respectively, to the resulting toxin gene set that we downloaded from the Pfam database (33) in June 2021 and manually curated based on their functional description. We mapped these domains to genes from the IMG genome database using HMMER3.0 as described before (34). Altogether we found additional 895,927 proteins coding genes that include the toxin or the immunity protein domains. The resulting dataset was then manually curated for quality assurance, and toxin or antitoxin genes erroneously included were removed.

**Figure 1.**
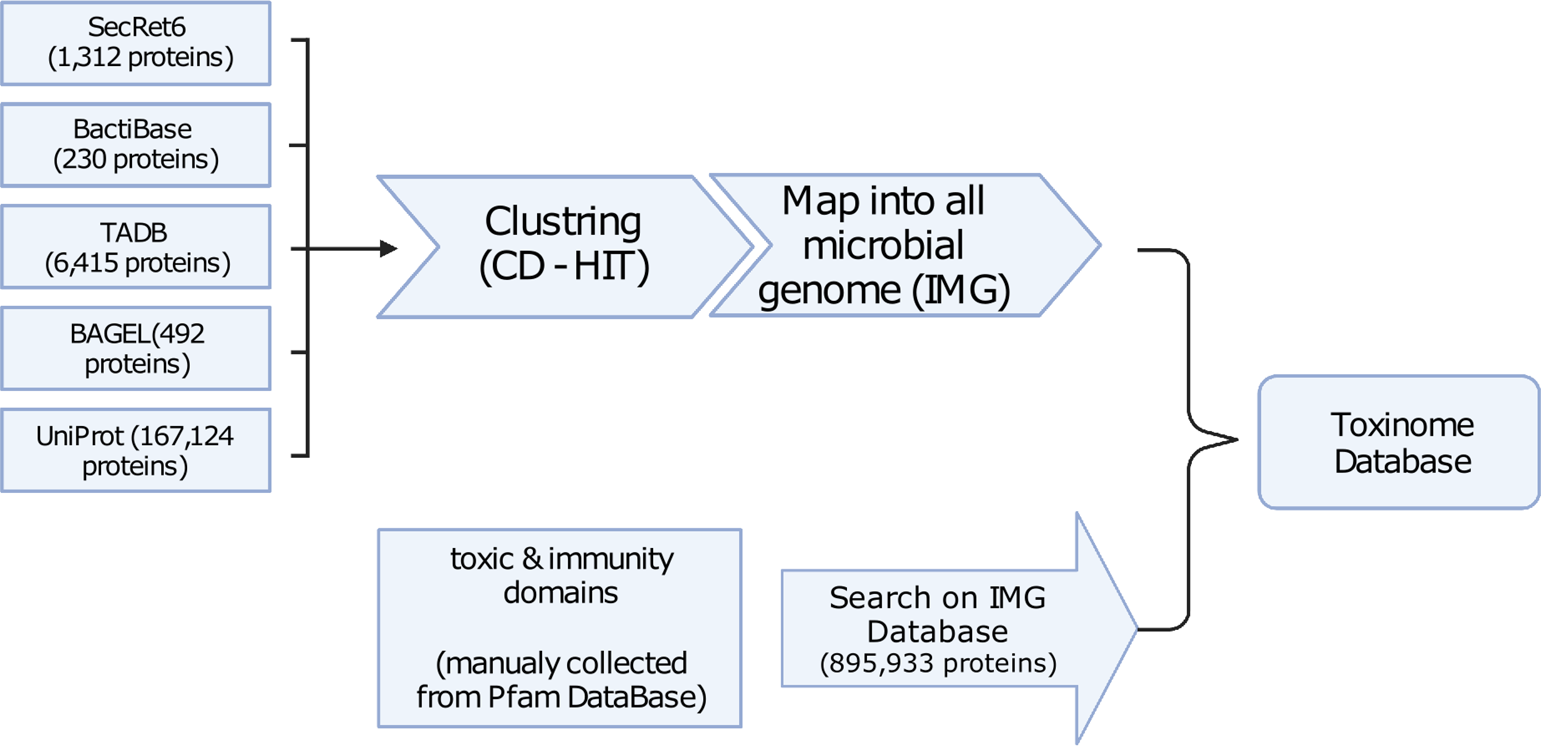
Construction of Toxinome database. The toxin proteins from Secret6, BactiBase, TADB, Bagel, and UniProt databases were joined, clustered to remove redundancy, and were mapped to microbial genomes from Integrated Microbial Genomics (IMG) database. In addition, we manually curated and retrieved toxin and immunity domains from Pfam, and map them again into proteins and genes from IMG. The union of these sets resulted in Toxinome.

### Toxin Island Prediction

We systematically searched DNA scaffolds from bacterial genomes from the Integrated Microbial Genomes (IMG) database for segments where toxins and antitoxins from the toxinome were closely located to each other. Different values for the parameter that represents the maximum distance between two adjacent toxins or antitoxins were tested, ranging from 10Kb base pairs (bp) to 60Kb bp. This analysis generated multiple DNA segments that are rich in toxins and antitoxins.

We employed two scoring methods to assess the segments, one based on the Poisson distribution and another utilizing a specialized scoring approach. To evaluate the background density of toxinome genes within genome segments, we assumed a Poisson distribution: 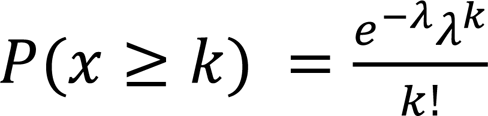. This involved calculating the overall presence of toxins and antitoxins per unit length in the genome, which was then multiplied by the segment’s length to calculate the expected value of lambda. K represents the actual number of toxins and antitoxins within the segment. This analysis generated a p-value, which quantifies the probability of observing a grouping of toxins and antitoxins in the genome that is at least as dense as the one observed. Additionally, we assigned a score to each hit, considering factors such as the number of toxins and antitoxins (T), the total gene count (G), and the length of the segment (L). Several parameter combinations were tested to establish an accurate measure of a Toxin Island, and after careful evaluation, the most optimal final formula was determined to be 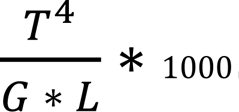. A high score represents a DNA segment enriched with toxins and antitoxins.

After scoring all the segments, we ensured the reliability of our findings by establishing thresholds for characterizing Toxin Islands: a score of > 4, a Poisson p-value ≤ 0.05, and a minimum of four toxins within a segment. The threshold of a score > 4 was chosen as it showed the best improvement in the average Poisson p-value of our hits (Supplementary Figure 1). This process resulted in 23,025 defined as “Toxin Islands”. Subsequently, we eliminated redundant results using mmseqs2 (35) (min-seq-id=0.99, seed-sub-mat=3, cluster-mode=2) and chose the best scoring representative, to obtain a final set of 5,161 unique “Toxin Islands” derived from 4,240 distinct bacterial genomes.

### Annotation of hypothetical proteins as Toxins and Antitoxins

We employed a two-step approach to predict the structures and functionally characterize unannotated proteins within the Toxin Islands. Firstly, we utilized ColabFold (36), a deep learning-based method, to predict the structures of these proteins. ColabFold utilizes the PDB file ranking system, and we selected the PDB file with the highest rank from the output files.

Subsequently, for the functional characterization of these proteins, we employed Foldseek (37) as our tool of choice. Foldseek enables structural alignment and identification of similar proteins, providing valuable insights into their structural characteristics and potential functional roles. To achieve this, we conducted a search of the proteins against the AlphaFold/UniProt50 v4 and PDB100 2201222 databases, utilizing Foldseek’s built-in search functionality. To ensure the selection of highly relevant and reliable matches, we implemented specific thresholds for filtering the results obtained from Foldseek. The applied thresholds included an e-value lower than 0.003, a TM-score higher than 0.6, and a minimum coverage of at least 90% of the query protein.

### Design of the Toxinome website

The Toxinome website was constructed using the Django framework (https://www.djangoproject.com) for full stack web development in Python. SQLite database engine was used for storing the collected data, and DIAMOND software (38) was used for online homology-based search, as part of the backend development of the Toxinome website. The data are organized in five SQL tables - genome table, gene table, cluster table, Pfam domain table, and Pfams in genes table. Each table stores the relevant information about collected toxins and antitoxins. The tables have associated connections based on common fields, which enables retrieval of different intersections of data and their presentation with simple frontend code. HTML, CSS, JAVA SCRIPT and bootstrap toolkit (https://getbootstrap.com/) were used for frontend development. Finally the constructed website was deployed on the pythonanywhere server and can be accessed by URL: http://toxinome.pythonanywhere.com. The website was tested and found working from chrome, internet explorer, firefox and safari browsers.

### Phylogenetic analysis

In order to build a phylogenetic tree to represent toxin and antitoxin prevalence across bacterial classes, we chose a random representative genome from each taxonomic class of our dataset and used these to construct a tree in which each leaf represents a taxonomic class. As a basis for comparison, we used universal marker genes. Specifically, we used 29 COGs out of 102 COGs that correspond to ribosomal proteins= (39). The COGs used are: COG0048, COG0049, COG0051, COG0052, COG0080, COG0081, COG0087, COG0088, COG0089, COG0090, COG0091, COG0092, COG0093, COG0094, COG0096, COG0097, COG0098, COG0099, COG0100, COG0102, COG0103, COG0185, COG0186, COG0197, COG0198, COG0200. COG0244, COG0256, COG0522. We aligned each COG protein from each representative genome using Clustal Omega version 1.2.4 (40). We then concatenated all the COG protein alignments for each genome, filling missing COGs with gaps. We used this 29 COG protein (single copy marker protein) concatenated alignment as an input to FastTree2 with default parameters (41) to create the phylogenetic tree. R’s ggtree v2.4.2 (42) and ggtreeExtra v1.0.4 (43) packages were used to plot the tree.

### Toxin and antitoxin correlation with microbial environment temperature

Metadata of temperature range (mesophile, thermophile, hyperthermophile, psychrophile) of different microbes that are included in Toxinome were downloaded from IMG database. For each genome we also calculated the proportion of toxin and antitoxin genes as part of the total gene number.

## Results

### Toxinome content and website usage

We downloaded 175,573 toxin sequences of various types from five databases, with Uniprot being the main one, and mapped them into 59,475 bacterial genomes to find all their orthologous and paralogous proteins (Materials and Methods). In addition, we manually curated toxin and immunity (antitoxin) protein domains and mapped them to genes in our genome dataset (Materials and Methods). This process resulted in the Toxinome database. The proteins were further clustered into 70,867 clusters (threshold = 0.7, alignment coverage = 0.92) using CD-HIT software . Next we developed a user-friendly website, http://toxinome.pythonanywhere.com, to store the toxin information online, and facilitate querying our toxin/antitoxin database (Figure 2). When browsing by organism, the user can find an alphabetically ordered list of all toxins and antitoxins from a selected organism. The information is displayed in a tabular way, including the Integrated Microbial Genomics (IMG) gene ID (44), product (protein) name, DNA scaffold, gene positions on the scaffold, protein length in amino acids, Pfam domains within the protein, functionality (Toxin/Antitoxin), information source, and cluster identifier. By clicking the Pfam link on the organism information page, the website redirects to a page describing the Pfam domains page that are encoded within the gene. Each Pfam has its Pfam ID, Pfam name, domain classification (Tox/Anti-Tox), coordinates on the protein, length in amino acids, and its graphic representation. Pfam ID is an active link to the record on the UniProt website (45). The website redirects to the cluster page by clicking on the cluster number link on the organism information page. The cluster page shows all the genes in the specific cluster with their information. The user can retrieve all the genes that have a specific Pfam name by choosing the Pfam name from the Pfam name list under the “Browse By Pfam’’ button. “Perform advanced Search*”* functionality allows searching for records using free language search for one of the four most usable fields: product name, organism name, Pfam name, Pfam ID. The combination of these fields can also be used to retrieve the records that contain the intersection of the searched fields. “Homologous Protein Search*”* allows a search based on an amino acid sequence similarity based on a query protein sequence. The amino acid query sequence is aligned to the protein database using DIAMOND (32) with the following parameters: query cover of 90%, reference protein cover of 60%, minimum identity of 40% and e-value of 0.001. The best 25 hits are presented with all their genetic information. Finally, the user can download the entire database as a comma-separated file with all the database information, a FASTA file with all the protein sequences, and Toxin Island information.

**Figure 2.**
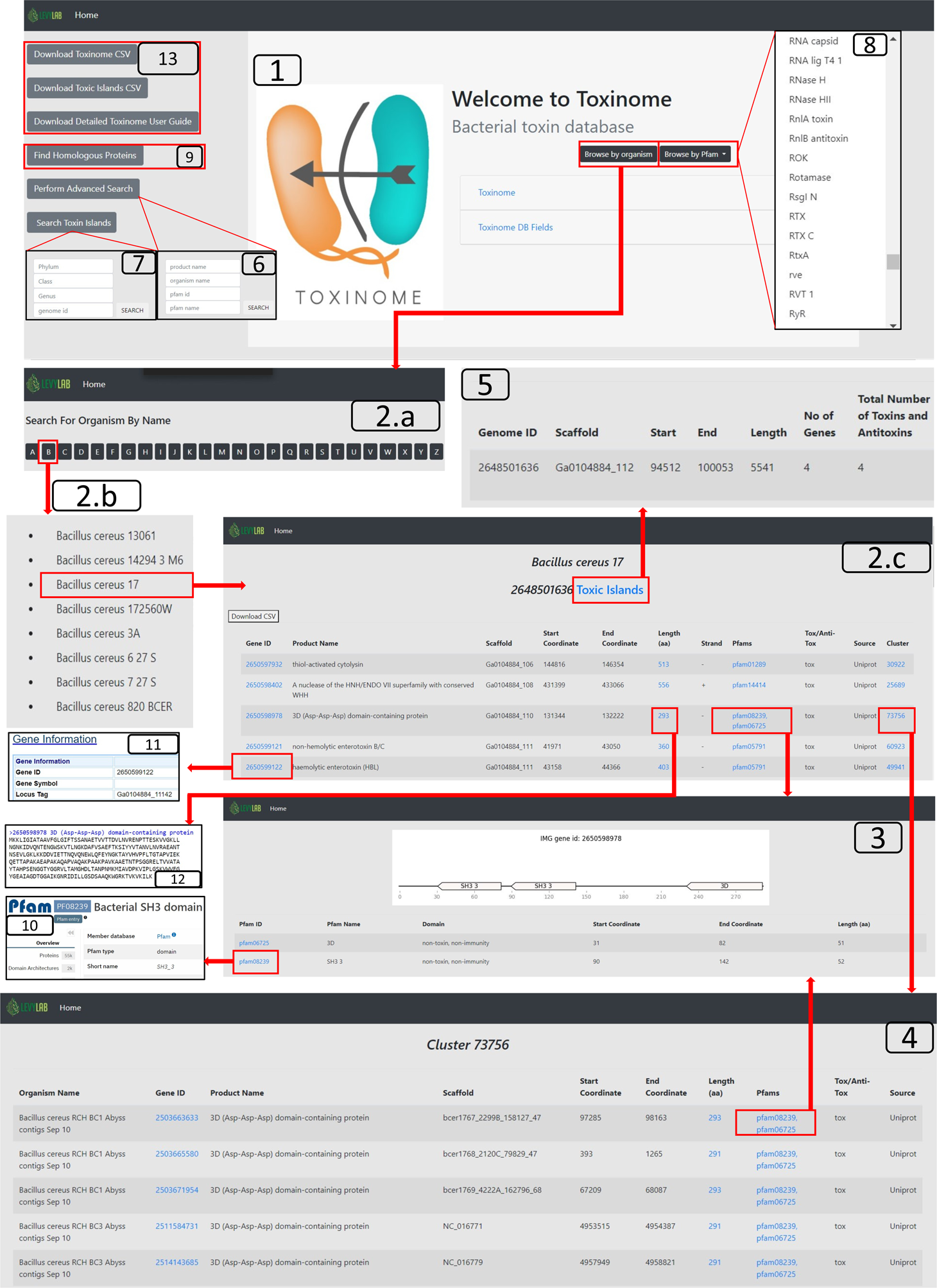
Schematic representation of the functionalities available on the Toxinome. The homepage [1] serves as the starting point for accessing the database.The user can browse the database by organism names organized in alphabetical order [2.a]. Selecting an organism name from the list [2.b] provides information on toxins and immunity genes encoded in the genome [2.c]. Each protein is associated with internal Pfam domains [3], and genes are part of protein clusters [4]. The database provides access to the toxin islands of the organism [5]. A more advanced search [6] allows users to search for a protein domain of interest. Advanced queries for toxin islands can be made based on Phylum, Class, Genus, and IMG genome id [7]. The database can be filtered by pfam name using a list of existing domains [8]. Sequence-based search [9] enables the detection of homologous sequences. Links associated with pfam id and gene id on an organism page redirect to external databases (https://www.ebi.ac.uk/interpro) [10] and (https://img.jgi.doe.gov/) [11]. Clicking on the length link provides access to the protein sequence on the IMG website [12]. The user can download the entire database, the toxin islands table, and the detailed user guide [13].

### Distribution of toxin and antitoxin across prokaryotes

We analyzed the toxin and antitoxin content of bacteria and archaea by calculating the average toxin gene per genome in class (number of toxins in class / number of genomes in class). The results are presented on a phylogenetic tree that was constructed based on single copy marker genes (Materials and Methods). As we expect, there is a high correlation between toxin and anti-toxin content (R = 0.6581, p-value = 7.59×10^−13^). The classes that are highly rich with toxins are Actinobacteria (Actinobacteria phylum), Gloeobacteria (Cyanobacteria), Caldilineae (Chloroflexota), Gammaproteobacteria and Acidithiobacillia (Pseudomonadota, previously Proteobacteria). Compared with bacteria, Archaeal classes (“hours” 3-4 in Figure 3) have relatively low content of toxins and antitoxins. Interestingly, we noted that bacterial classes that are depleted for toxins and antitoxins are surprisingly enriched in many thermophilic and hyperthermophyilic classes from eight different bacterial phyla. These classes include Aquificae (Aquificota phylum), Thermotoage (Thermotogota phylum), Dictyoglomia (Dictyoglomota phylum), Caldisericia (Caldisericota phylum), Thermomicrobia (Thermomicrobiota phylum), Calditrichae (Calditrichota phylum), Coprothermobacteria (Coprothermobacterota phylum), and Methylacidiphylae (Verrucomicrobiota phylum). The Caldilineae thermophilic class is an exception of being toxin-rich. Other classes that are depleted in toxins and antitoxins are Dehalococcoidia, Endomicrobia, Cathonomonadetes. Chitinivibirionia, Chlamydia, and Deferribacteres.

**Figure 3.**
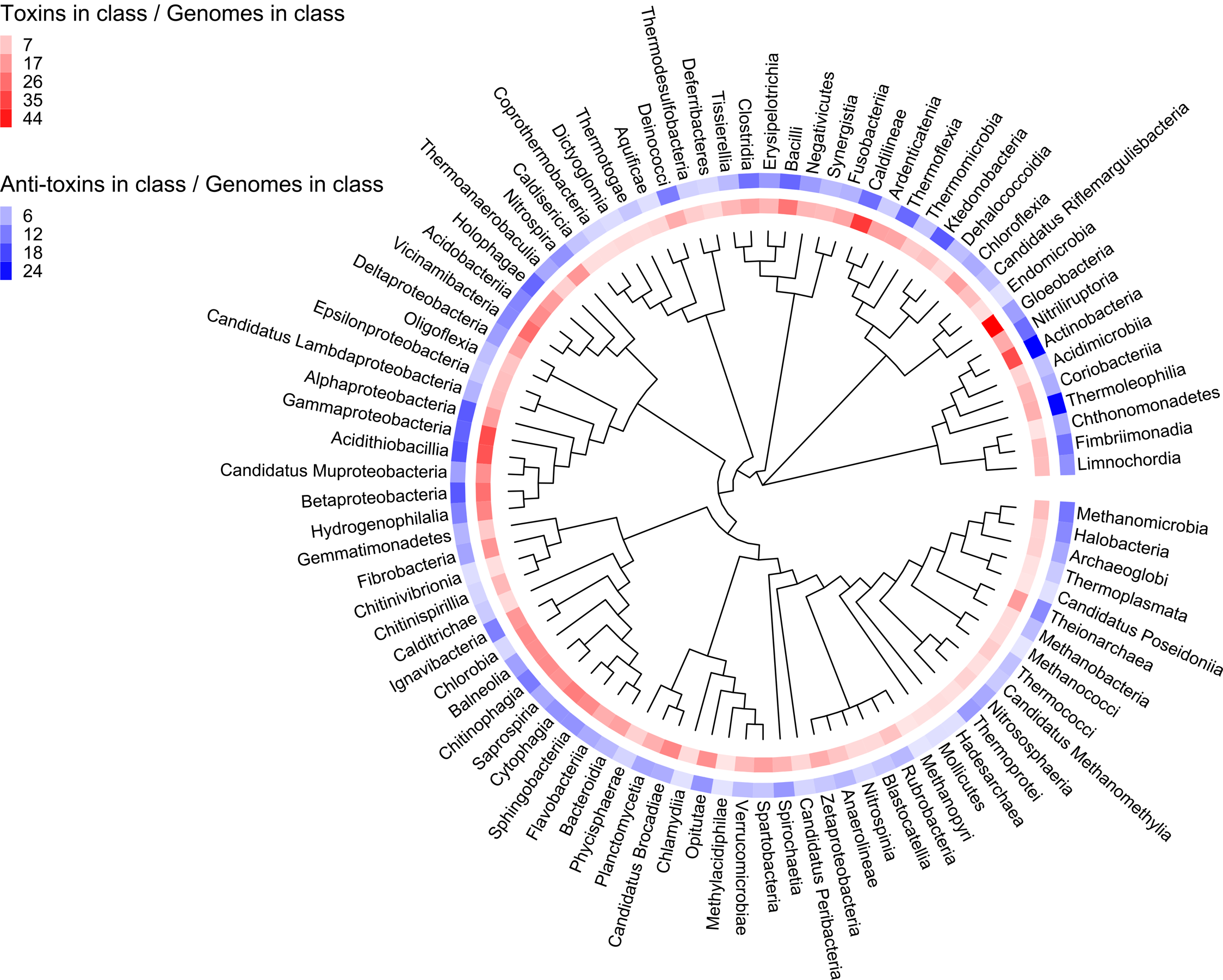
Toxin and Antitoxin content in bacterial and archaeal genomes. Toxinome genes were counted in each class and were divided by the number of genomes per class.

To further test if toxin and antitoxin gene frequency is correlated with bacterial adaptation to temperature we used 11,748 publicly available bacterial and archaeal genomes (44) for which the temperature preference metadata is available, namely whether the microbes are mesophiles, thermophiles or psychrophiles. We found that mesophiles have significantly higher content of toxins and antitoxins than thermophiles and psychrophiles bacteria. Statistical significance was calculated by ANOVA test followed by pairwise comparisons and adjustment with Tukey method. The results are summarized in Tables 1-4.

**Table 1.**
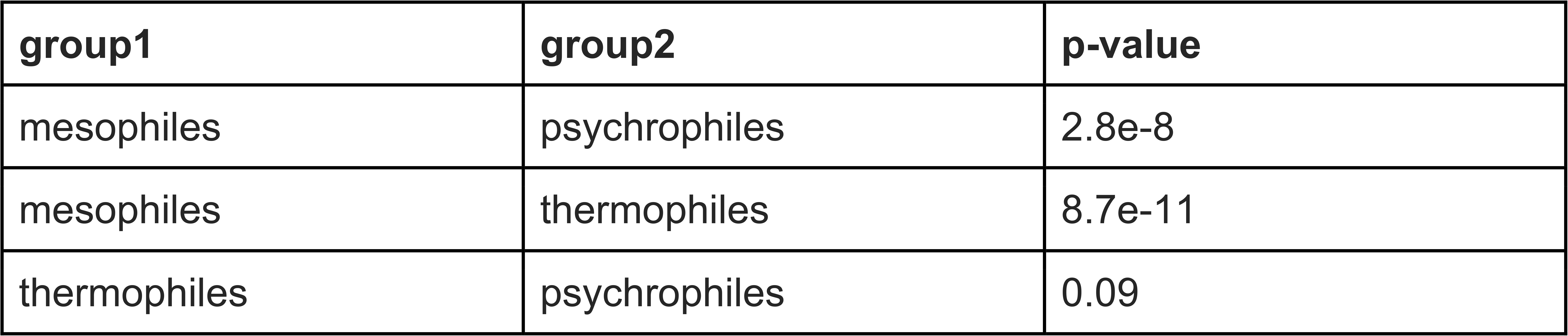
Distribution of Bacterial Antitoxins.

**Table 2.**
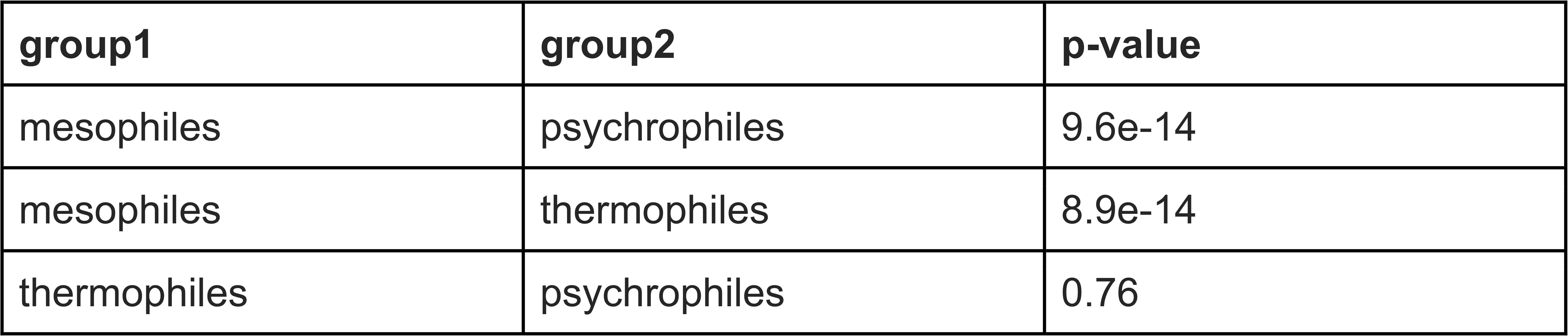
Distribution of Bacterial Toxins.

**Table 3.**
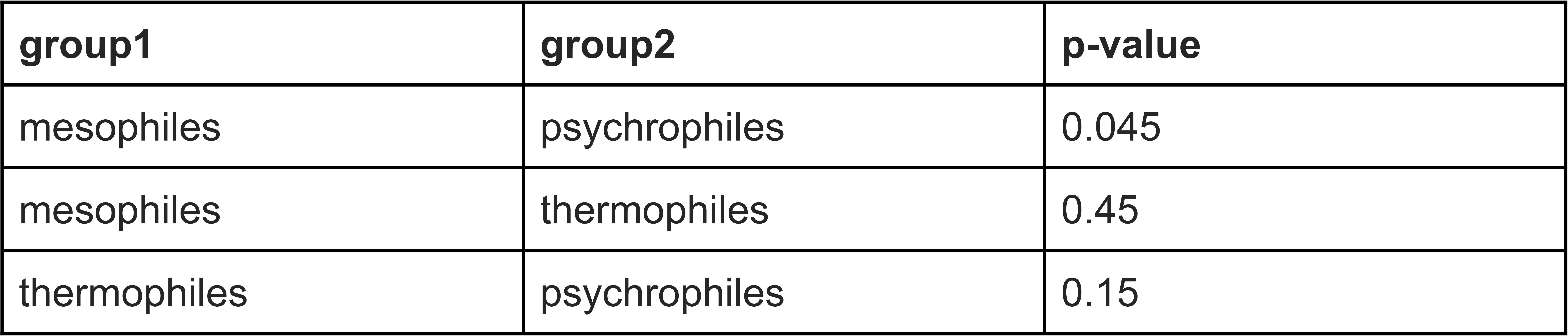
Distribution of Archaeal Antitoxins.

**Table 4.**
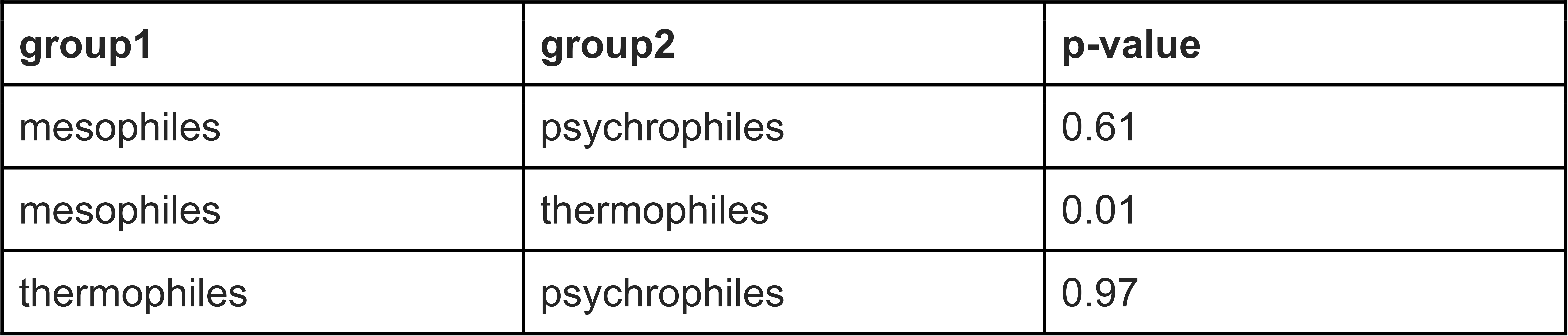
Distribution of Archaeal Toxins.

As can be seen from the above, the bacterial mesophiles have significantly more toxins and antitoxins than bacterial psychrophiles and thermophiles when normalized to their genome size (Figure 4). For archaea the difference between mesophiles and psychrophiles groups has borderline p-value in both toxins (p-value ≤ 0.0138) and antitoxins (p-value ≤ 0.0455) count (Figure 4)

**Figure 4.**
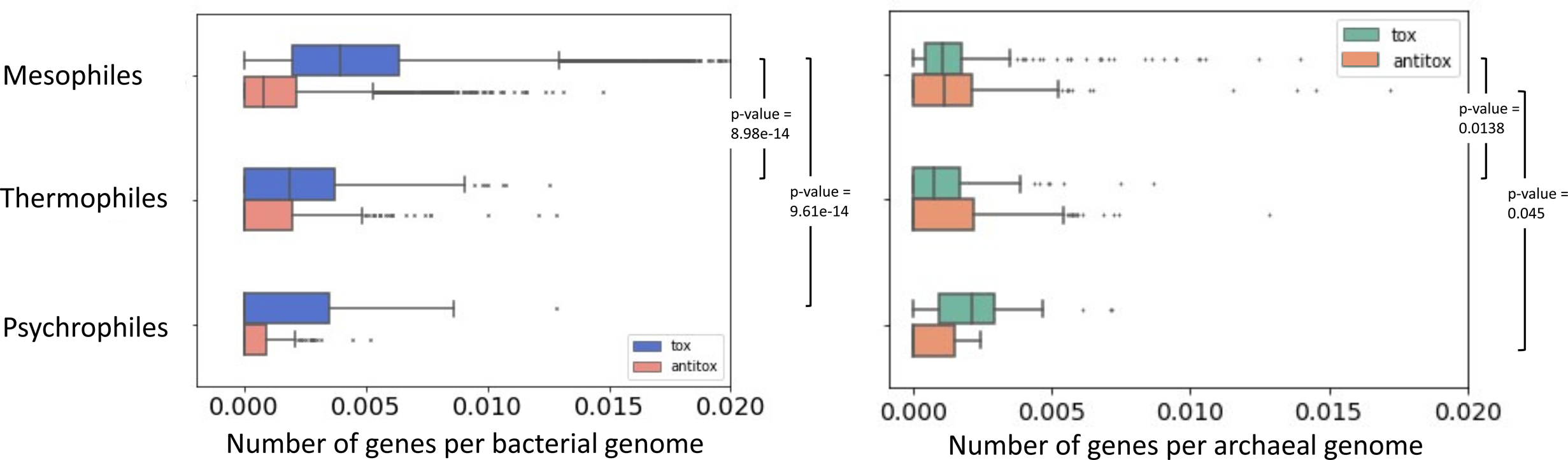
Toxin and antitoxin genes are enriched in mesophilic bacteria. Toxin and antitoxin counts were normalized by genome size in bacterial and archaeal genomes.

### Identification of *Toxin Islands*

Genomic pathogenicity islands were described as genomic regions of 10-200Kb that are enriched in virulence genes, such as toxins, adhesins, invasins, iron uptake systems, and protein secretion systems, and are enriched in pathogens (46–49). These regions often have GC content different from the genome, are rich with mobile genetic elements and repetitive sequences, and are frequently horizontally transferred and integrated next to tRNA genes (46, 50). *Erwinia Amylovora*, for instance, has been reported to have a number of pathogenic islands, mostly those associated with the type II and III secretion systems (T2SS and T3SS) (51). *Salmonella* genus contains multiple pathogenicity islands which encode T3SS and its effector genes, and genes required for survival inside macrophages (52, 53), and the pathogenicity islands of *Pseudomonas aeruginosa* are required for virulence in animals and plants (54). A few years ago Bacteroidales species in the human gut were reported to encode dynamic genomic islands of immunity genes that protect against T6SS antibacterial effectors of co-resident microorganisms (55).

Our observation of toxins and antitoxins frequently grouped together led us to propose the existence of “Toxin Islands” which are genomic islands enriched in toxins and antitoxins within bacterial species. We hypothesized that some pathogenicity islands are enriched in toxin genes and therefore would be more accurately functionally annotated as “Toxin Islands”. In addition, Toxin Islands can have non-virulence function when they overlap with antiphage defense islands which are rich with toxin-antitoxin genes (56, 57) or enriched in antibacterial arsenal used for outcompeting microbes.

To test this hypothesis, we developed a statistical, score-based, computational algorithm capable of identifying regions in a genome that exhibit high densities of toxins and antitoxins. The algorithm systematically scans each genome, searching for segments where adjacent toxins or antitoxins are within a specified maximum distance threshold (Materials and Methods). Subsequently, we applied stringent filtering using carefully selected thresholds to delineate the regions that meet the criteria for classification as “Toxin Islands” (Material and Methods, Supplementary Figure 1). Using this method, we identified 23,025 Toxin Islands where 5,161 of them are unique islands that originated from 4,240 different genomes (Supplementary Table 2). The majority of the islands have lengths shorter than 15Kb (Supplementary Figure 2), indicating a prevalence of relatively compact Toxin Islands. The phylogenetic origin of these Toxin Islands is highly diverse and spans across 50 different classes. However, they are most abundant in the classes of Gammaproteobacteria, Bacilli, and Actinobacteria. (Figure 5).

**Figure 5.**
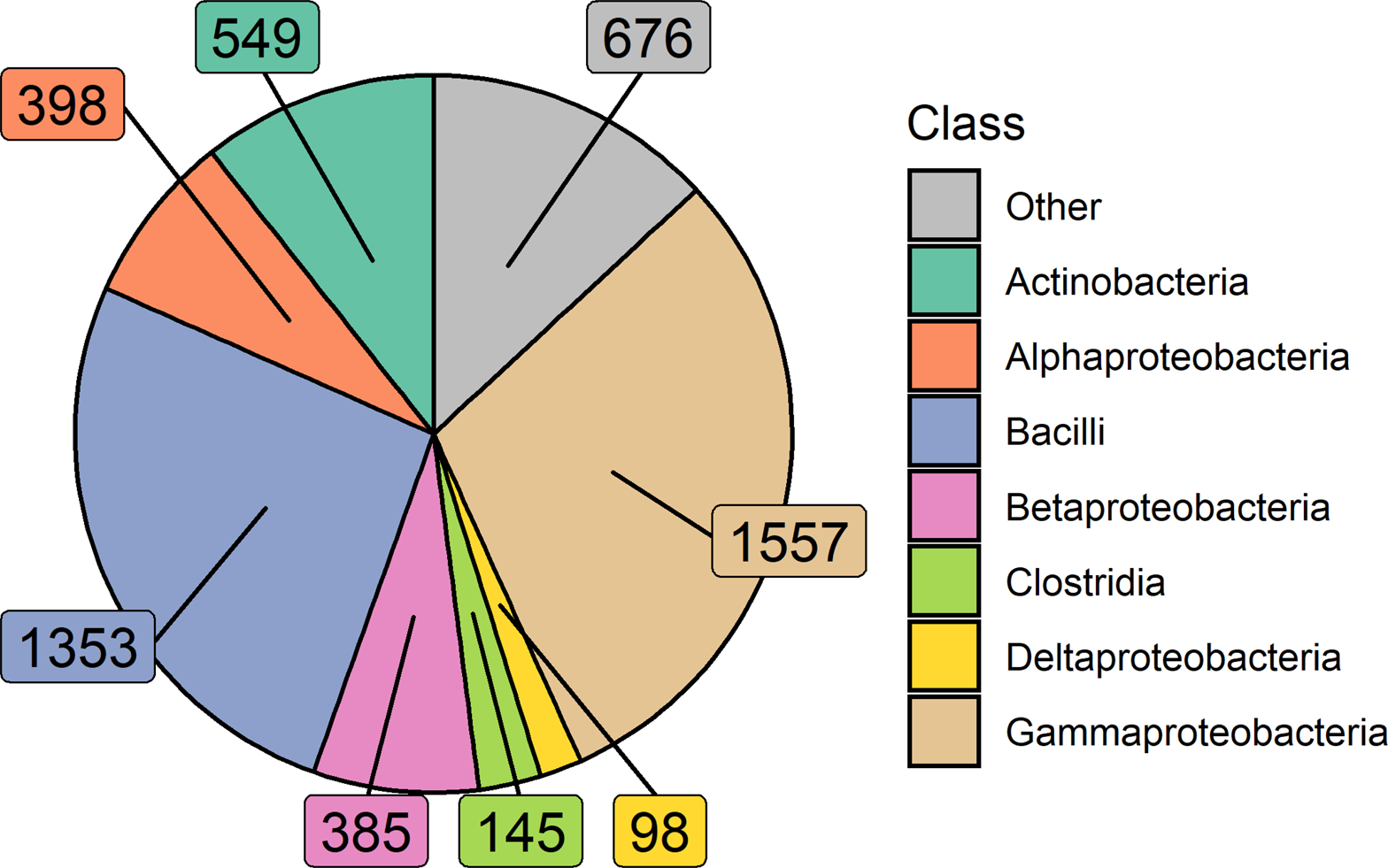
Number of unique Toxin Islands identified per class. This plot illustrates the distribution of unique Toxin Islands discovered across different classes. Each slice represents a specific class, and the size of each slice corresponds to the number of distinct Toxin Islands identified within that class. The term “Other” refers to classes that either exhibit low toxin island counts or genomes that lack taxonomy annotation.

To characterize the identified Toxin Islands across different classes, we conducted a count of toxins and antitoxins for each of the abundant classes in our analysis (Figure 6). Interestingly, we observed a consistent trend where the majority of Toxin Islands contained a higher number of toxins compared to antitoxins. This trend was particularly pronounced in the class Bacilli, where over half of the identified islands exclusively contained toxins without any associated antitoxins. Furthermore, we found that Toxin Islands often tend to contain homologous toxins or antitoxins within the same island, suggesting the occurrence of local gene duplication events (Supplementary Figure 3).

**Figure 6.**
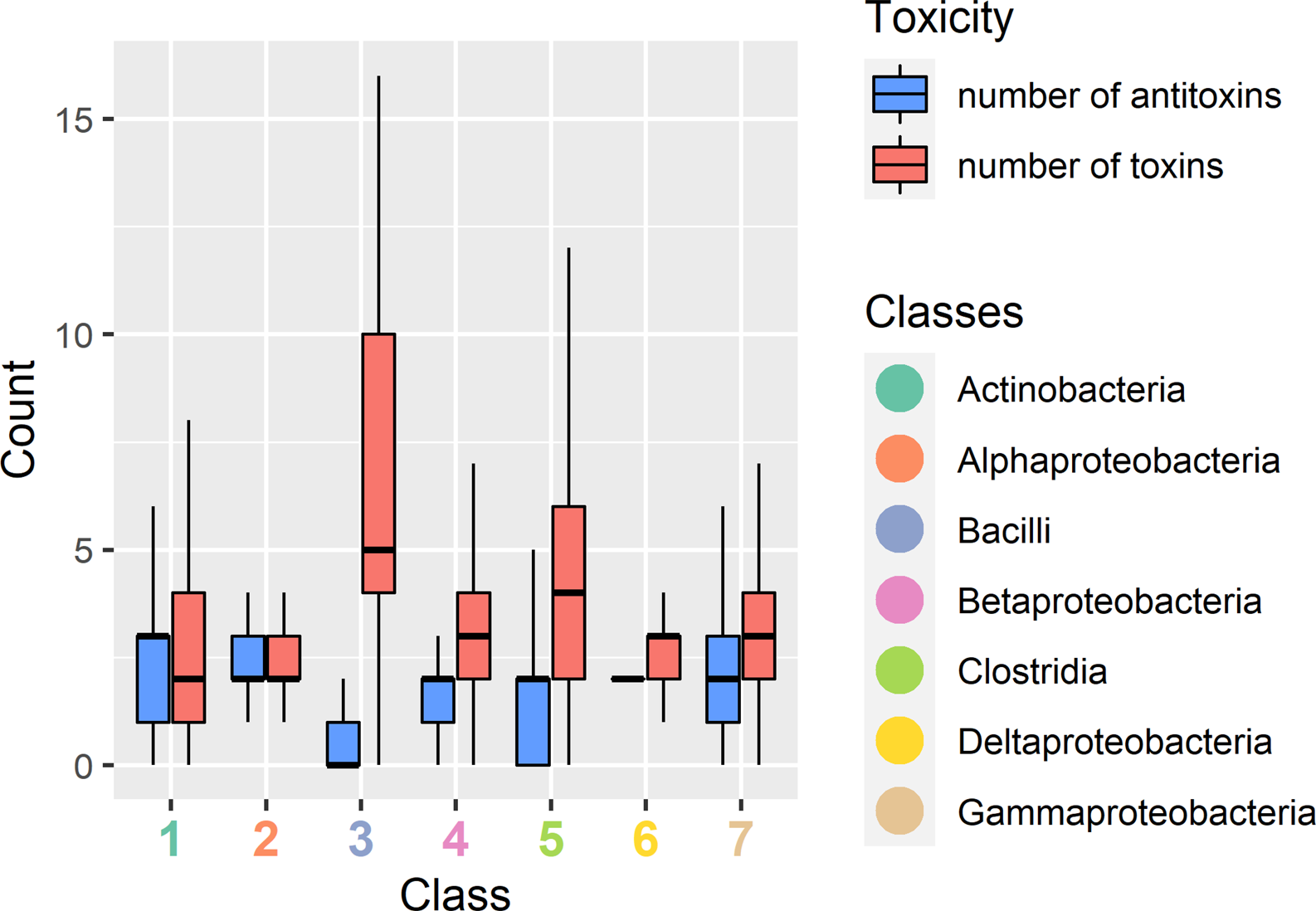
Toxin and Antitoxin Counts in Toxin Islands per Class. The plot represents the distribution of toxins and antitoxins within each identified Toxin Island. The number of toxins is depicted by the red box, while the number of antitoxins is represented by the blue box. The Y-axis indicates the count of toxins or antitoxins, while the X-axis corresponds to the classes that are rich in Toxin Islands. Each number on the X-axis corresponds to a specific class as follows: 1=Actinobacteria, 2=Alphaproteobacteria, 3=Bacilli, 4=Betaproteobacteria, 5=Clostridia, 6=Deltaproteobacteria, 7=Gammaproteobacteria.

One example of a predicted Toxin Island is found in the denitrifying bacteria *Thauera phenylacetica B4P* (Figure 7A) (58). This specific island spans a length of 15,000 base pairs and contains 9 toxins and 9 antitoxins. Notably, within this Toxin Island, among other toxins and antitoxins, one can identify the presence of the toxin YoeB accompanied by the antitoxin YefM (Figure 7A). In addition to the toxins and antitoxins, other genes present in this island include genes that encode to type IV pilus assembly protein, Transposase, Glycosyltransferase, and hypothetical proteins (Figure 7A).

**Figure 7.**
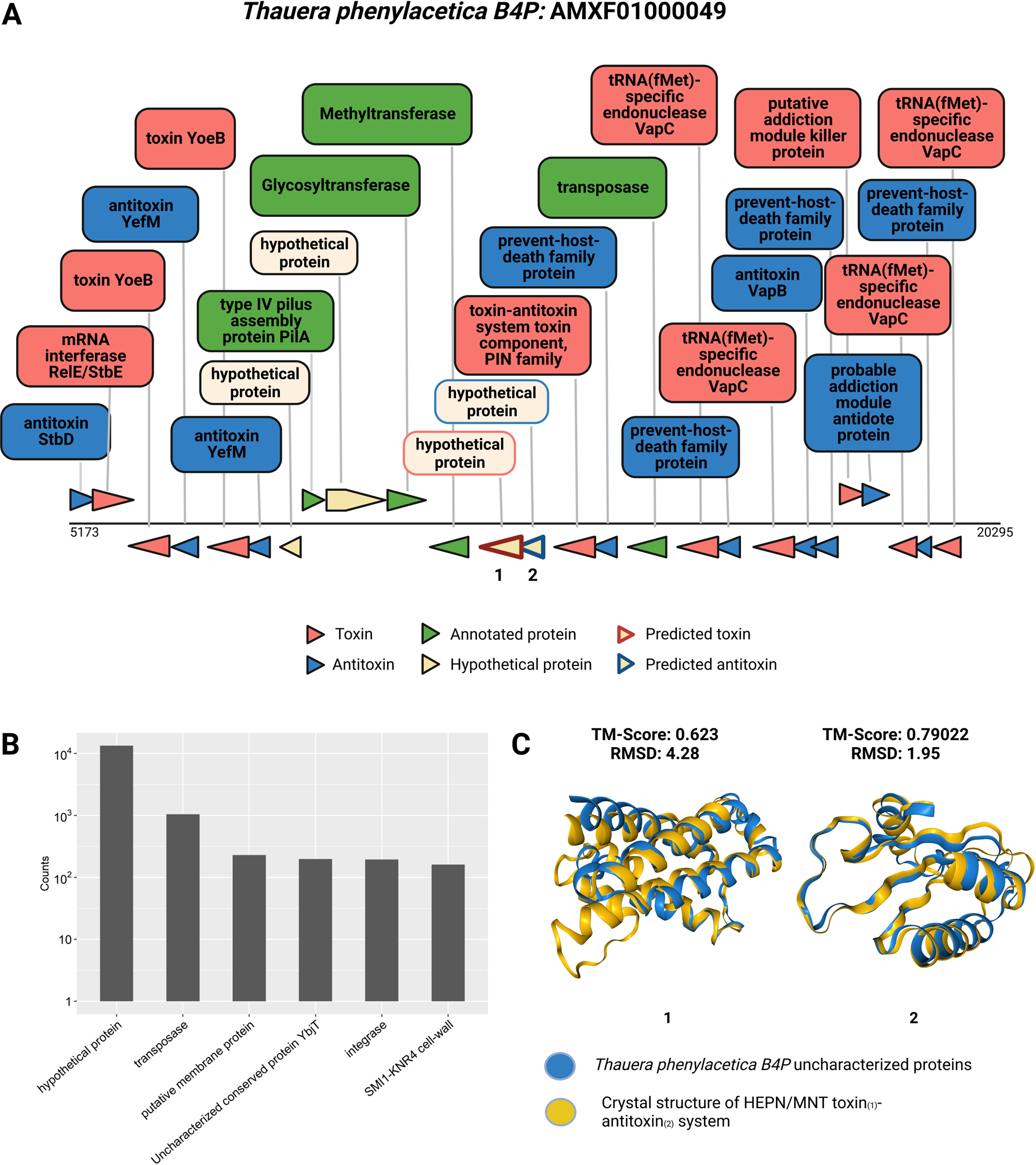
Toxin Islands occasionally encode unidentified toxins and antitoxins. **A. Identified Toxin Island within *Thauera phenylacetica B4P***. The figure displays a Toxin Island identified within the genome of *Thauera phenylacetica B4P,* highlighting specific proteins and their respective functions. Different colors are used to distinguish toxins, antitoxins, functionally annotated proteins, and hypothetical proteins. Proteins for which we performed computational functional characterization are indicated by colored frames. The corresponding structures for these characterized proteins are shown in Figure 7C, labeled as 1 and 2. **B. Distribution of protein annotations in Toxin Islands for non-toxin/non-immunity genes.** Each bar represents a frequently observed product annotation associated with genes found in Toxin Islands, excluding toxins and antitoxins. The y-axis represents the counts presented in logarithmic scale. **C. Protein structure alignment of uncharacterized proteins within the toxin island:** Uncharacterized proteins are colored in blue, while known proteins from the aligned database are colored in yellow. Protein numbers 1 and 2 displayed similarity to the crystal structure of HEPN and MNT toxin-antitoxin system (PDB Structures ID: 7AE8 and 7BXO) respectively, identified in *Shewanella oneidensis MR-1*.

We hypothesized that toxins or antitoxin genes can be identified by analyzing proteins with unknown activity that are densely clustered in Toxin Islands. In our analysis of non-toxin and non-antitoxin proteins within these islands, we observed that a significant proportion of protein coding genes lack any Pfam domain annotation and are annotated as “hypothetical proteins” (Figure 7B). These genes may potentially encode novel toxins and antitoxins, which genomically cluster together with the known toxins and antitoxins. To test this suggestion, we investigated the potential functions of the unannotated proteins within the specific island depicted in Figure 7A, and employed a comparative analysis approach. Using Foldseek (37), we conducted a structural comparison of these proteins against protein structure databases. Remarkably, we found that two hypothetical protein-coding genes, labeled as 1 and 2, displayed a high degree of similarity to known toxin and antitoxin systems (Figure 7C). Specifically, protein numbers 1 and 2 exhibited similarity to the crystal structure of HEPN/MNT toxin-antitoxin system found in *Shewanella oneidensis MR-1*. The MntA antitoxin (MNT-domain protein) acts as an adenylyltransferase and chemically modifies the HepT toxin (HEPN-domain protein) to block its toxicity as an RNase (59). These findings provide compelling evidence supporting the hypothesis of the presence of novel toxin and antitoxin genes within these islands, highlighting the genomic clustering phenomenon. Another group of genes which is abundant within Toxin Islands are transposases and integrases (Figure 7B) which indicates that the islands can be transferred horizontally between bacteria.

## Discussion

Toxinome is a unique and extensive database of most known toxins and antitoxins in 59,475 bacterial genomes. The toxins include classic exotoxins such as the Anthrax and Pertussis toxins, toxin effectors of different secretion systems, toxin-antitoxin systems, bacteriocins, and others, along with their cognate antitoxins. We expect that the database will serve as a valuable resource for the large research community interested in the biology of bacterial protein toxins that are used for virulence, abortive phage infection, and inter-microbial competition. Toxinome database is presented through a user-friendly web interface that is easy to navigate, query, and download. A wide range of follow-up studies can be conducted using the data collected, from the discovery of new toxins, antitoxins, and toxin delivery systems and associating observed phenotypes with genes from Toxinome. Microbiologists can retrieve a full profile of all known toxins encoded by the microbe they study.

By analyzing the Toxinome database we identified prokaryotic classes that are rich and poor in toxins and antitoxins. It is intriguing to know whether the observed depletion in thermophilic and psychrophylic bacteria is the result of purifying selection against toxin and antitoxin gene presence in these clades. This can result from having less microbial competitors or less hosts for bacteria dwelling in extreme environments making toxins unnecessary energetically expensive molecular weapons. It was reported that when temperature increases, both microbial species diversity and Pfam diversity steadily decline, and hence this may also affect toxin and antitoxin gene content, particularly those used in inter-microbial competition in polymicrobial communities (60). We speculate that since most animals and plants dwell in mesophylic conditions there are less pathogens equipped with anti-eukaryotic toxins living in extreme temperatures. Alternatively, this finding might be the result of a bias in toxin and antitoxin functional annotation. Namely, it may suggest that most known toxins and antitoxins were discovered and studied in the organisms where they are supposedly enriched, such as human pathogenic proteobacteria, whereas biochemically poorly characterized clades may have independent sets of toxins and antitoxins that remain unidentified and can lead to description of new phenotypes in these microbes. Indeed, classic toxin-antitoxn systems were not studied in many extremophiles until quite recently (61) and novel toxin-antitoxin systems were predicted to be specific to thermophiles (15). It is also fascinating to test if the relatively low incidence of toxins and antitoxins is common to other extremophilic bacteria that live for example in acidic or alkaline environments. For example, certain hyperthermophilic and halophilic Archaea were described to produce bacteriocins called sulfolobicins and halocins, respectively (62).

Bacteria often tend to cluster together in the genome genes of similar functions such as antibiotic resistance (63, 64), heavy metal resistance (65), phage defense (56, 57), and pathogenicity (66). We utilized the Toxinome database to conduct an analysis and define the concept of genomic “Toxin Islands.” These islands exhibit a significant abundance of toxins and antitoxins within a short DNA stretch. We observed that the number of toxins surpasses the number of antitoxins in these islands, mostly in Bacilli, despite the fact that toxins and antitoxins are typically coupled together in the genome to safeguard bacteria from their own harmful weaponry. One possible explanation for this phenomenon is the presence of toxins that specifically target eukaryotes, rendering the bacteria immune to their own toxins. Alternatively, it could be due to the lack of annotation of antitoxins as Toxinome content is biased with threefold more toxins than antitoxins. In addition to their toxin and antitoxin content, these Toxin Islands are characterized by the presence of numerous genes, a majority of which have no functional annotation (Figure 7B). Moreover, these islands are abundant in mobile genetic elements, suggesting the horizontal transfer of genes into the islands or the lateral transfer of entire islands. It is plausible that the multitude of “hypothetical protein” genes found within Toxin Islands encode novel toxins and antitoxins that may target prokaryotic or eukaryotic cells, as we have shown in a Toxin Island form *Thauera phenylacetica* (Figure 7C). Consequently, annotating the Toxin Islands can greatly facilitate the discovery of gene functions, including those that lack sequence similarity to known genes, and play a pivotal role in shaping microbial interactions with hosts, phages, and other microbes. We anticipate that additional genes involved in toxin production, maturation, and secretion will also be localized in Toxin Islands. Exploring the cellular phenotypes resulting from the expression of hypothetical protein-encoding genes from the Toxin Islands of certain archaeal, thermophilic, and psychrophilic groups that we identified as having low toxin and antitoxin content could yield intriguing insights and novel types of toxins. Therefore, we believe that experimental testing of such toxins holds significant interest.

## Acknowledgements

AL is supported by the Israeli Science Foundation (Grants #1535/20, #3300/20, #3062/20), Alon Fellowship of the Israeli council of higher education, and the Volkswagen Foundation (Grant ZN 4041).

