## Supplementary Figures 1-3 for "Toxinome - The Bacterial Protein Toxin Database"

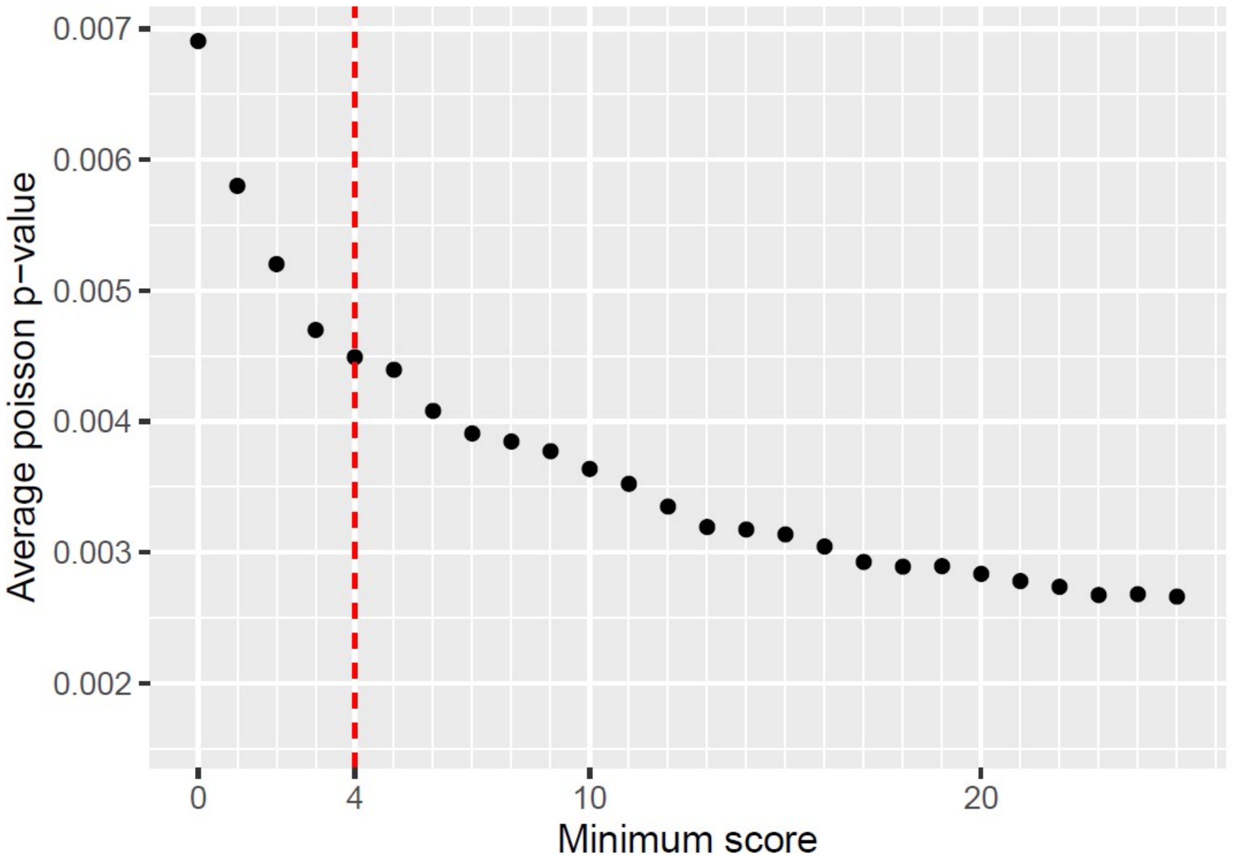


**Supplementary Figure 1. Average Poisson p-value of Toxin Islands in response to incrementally increasing the minimum score threshold:** The x-axis represents the minimum score cutoff of the results, and the y-axis represents the average Poisson p-value after the cutoff. The plot illustrates how the average Poisson p-value changes as the minimum score threshold increases. The selection of a minimum score threshold of 4 was made based on its superior improvement in the analysis.

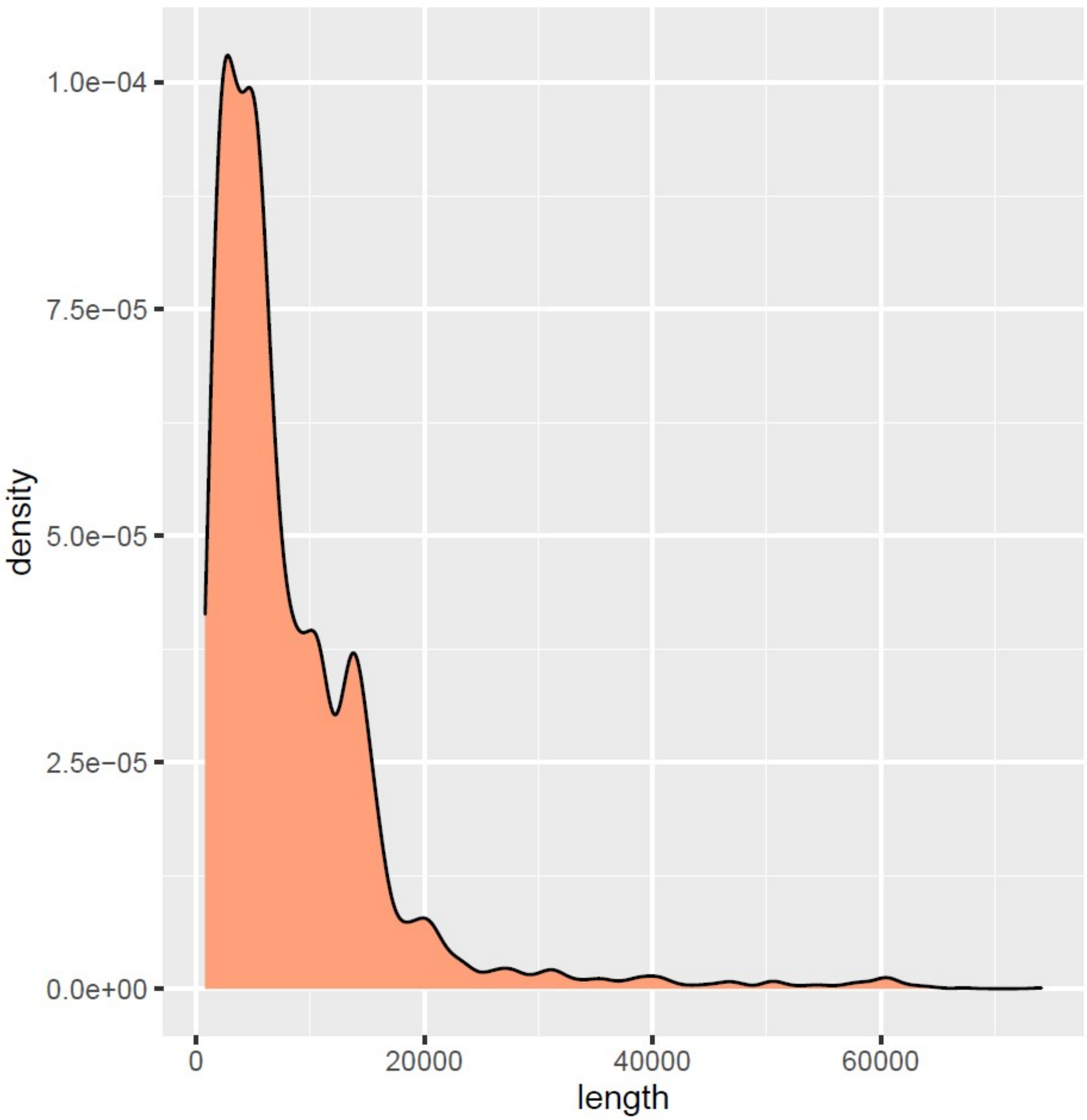


**Supplementary Figure 2. Density of Discovered Toxin Islands by Length.** This plot presents the density distribution of unique Toxin Islands' lengths in base pairs, as revealed by our analysis. The x-axis represents the length of the Toxin Islands, while the y-axis indicates the density or frequency of occurrence. It is observed that a majority of the islands have lengths shorter than 15 Kb, indicating a prevalence of relatively compact Toxin Islands.

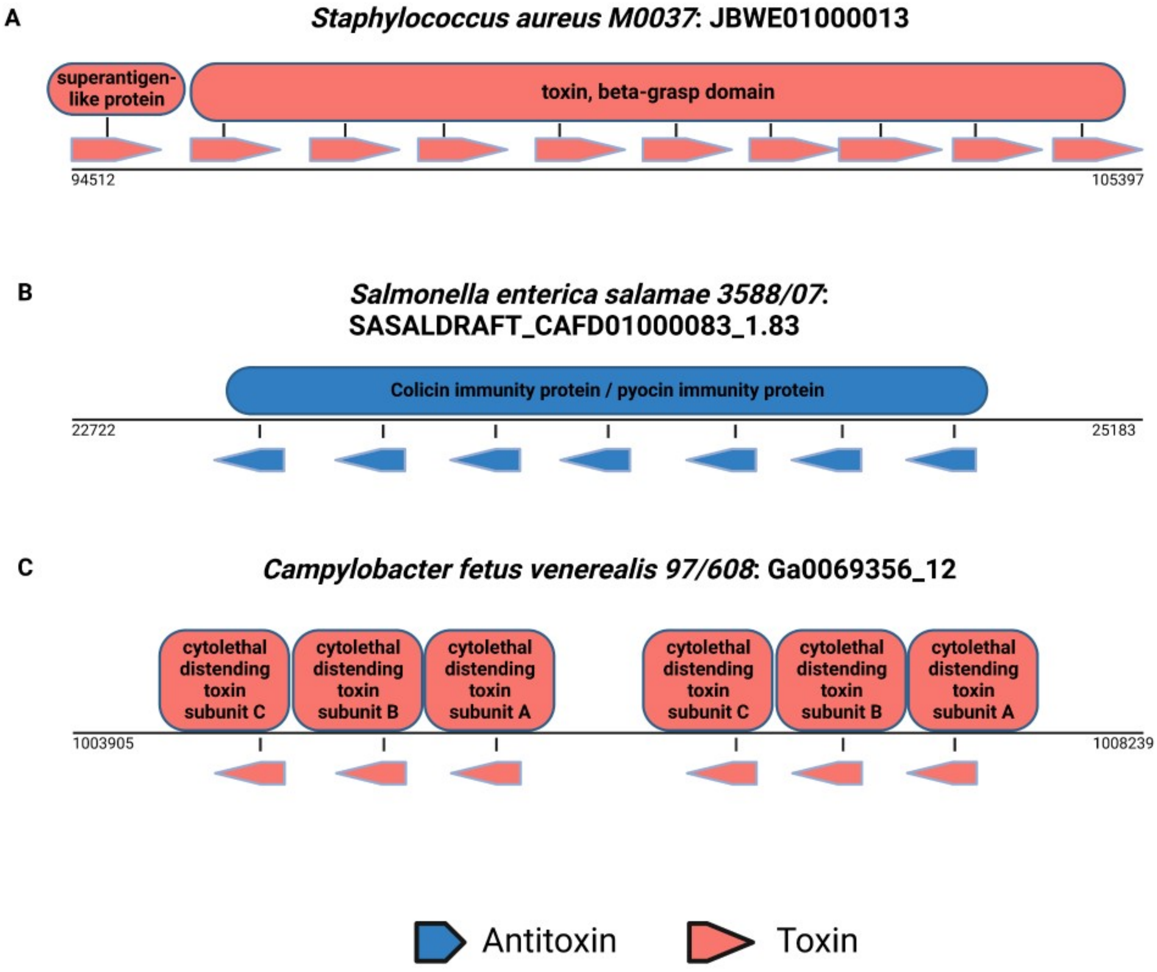


**Supplementary Figure 3 | Toxin Islands which contain homologous proteins.** The figure displays three distinct examples of Toxin Islands found in different species. In the illustration, red color indicates toxins, while blue color represents antitoxins. The function of each protein is described above its corresponding illustration on the genome. Proteins assigned with the same function description are considered homologous proteins, indicating their origin from a shared ancestor.
